# *CellPie*: a scalable spatial transcriptomics factor discovery method via joint non-negative matrix factorization

**DOI:** 10.1101/2023.09.29.560213

**Authors:** Sokratia Georgaka, William Geraint Morgans, Qian Zhao, Diego Sanchez Martinez, Amin Ali, Mohamed Ghafoor, Syed-Murtuza Baker, Robert Bristow, Mudassar Iqbal, Magnus Rattray

## Abstract

Spatially resolved transcriptomics has enabled the study of expression of genes within tissues while retaining their spatial identity. Most spatial transcriptomics technologies generate a matched histopathological image as part of the standard pipeline, providing morphological information that can complement the transcriptomics data. Here we present *CellPie*, a fast, unsupervised factor discovery method, based on joint non-negative matrix factorisation of spatial RNA transcripts and histological image features.*CellPie* employs the accelerated hierarchical least squares method to significantly reduce the computational time, enabling efficient application to high-dimensional spatial transcriptomics datasets. We assessed *CellPie* on two different human cancer types and spatial resolutions, showing an improved performance against published factorisation methods. Additionally, we applied *CellPie* to a highly resolved Visium HD dataset, demonstrating its high computational efficiency compared to standard non-negative matrix factorisation and other existing methods.

**Availability:** https://github.com/ManchesterBioinference/CellPie

## Introduction

In multicellular organisms, tissues are complex systems composed of millions of cells that constitute the building blocks of whole organs. Within tissues, cells vary in type and activity, and their development and function are influenced by interactions with their surroundings. Therefore, dissecting spatial cellular organisation and heterogeneity within tissues is important for understanding normal tissue function as well as diseases, which often have spatial origins (1).

Cutting-edge technologies, such as single-cell RNA sequencing (scRNA-seq), achieve high-throughput and high-resolution gene expression profiles, providing powerful insights into the characterisation of the heterogeneity at the transcriptomic level (2, 3). However, these methods dissociate the tissue, resulting in the loss of spatial dimension. In situ capture methods, such as spatial transcriptomics (ST) (4), commercially available as Visium by 10x Genomics, and Slide-seq (5) (Slide-seq V2), allow for molecular profiling while retaining the spatial information of the tissue, at various resolutions (6, 7). The barcoded Slide-seq beads (‘pucks’) provide near single-cell spatial resolution of 10*µm* while Visium barcoded spots have a coarser resolution of 55*µm*. Recently, 10x Genomics released the high-resolution Visium HD spatial gene expression platform (8), offering whole transcriptome spatial mapping at single-cell/near single-cell resolution (available at 2*µm*, 8*µm* and 16*µm*).

Spatial transcriptomics data offer a multi-modal view of the tissue landscape. In situ capture methods provide spatial and transcriptomic information along with matched Haematoxylin and Eosin (H&E) stained images as part of their standard pipelines. These histopathological images provide a comprehensive view of the tissue architecture that can complement spatial gene expression. Most current dimensionality reduction and factor analysis methods either model the gene expression modality alone or integrate together the spatial and gene expression parts, without utilizing the image modality. For example, dimensionality reduction methods such as the multi-modal NSF (and their Hybrid NSF - HNSF) (9) and MEFISTO (10), integrate spatial and gene expression dimensions, while unimodal methods, like probabilistic NMF (PNMF) and Factor Analysis (FA) (11) model gene expression alone. NSF is based on the non-negative spatial factorisation model which assigns an exponentiated Gaussian Process (GP) prior over the spatial locations with either a Poisson or negative binomial likelihood over the gene expression counts. MEFISTO performs a spatially aware factor analysis through factorising the gene expression data into latent GPs. While these methods scale reasonably well for Visium 10x datasets, when it come to higher resolved spatial transcriptomics datasets, such as the Visium HD, they become computationally prohibitive.

To address these limitations, we present *CellPie*, a flexible and scalable factor discovery tool based on unsupervised joint Non-negative Matrix Factorisation (jNMF). Both joint and single NMF-based methods have become popular in single-cell genomics for producing interpretable sparse features from high-dimensional data. Joint NMF has previously been applied to integrate multiple transcriptomics datasets (12), omics profiles from observational (TCGA) and experimental (CCLE) data (13), and biomarker dis-covery (14). *CellPie* jointly models spatial gene expression data and matched morphological imaging features, outputting a joint parts-based representation (factors) of the high-dimensional data in a simple and computationally efficient way. We benchmarked *CellPie* against published dimensionality reduction methods on two human cancer spatial transcriptomics datasets – a Visium 10x Genomics human prostate adenocarcinoma with invasive carcinoma and a Spatial Transcriptomics HER2-positive human breast cancer sample. Additionally, we applied *CellPie* to a Visium HD human colorectal cancer sample, demonstrating its high computational performance, against standard NMF implementations (15).

## Materials and methods

### Joint Non-negative Matrix Factorisation

*CellPie* employs a scalable NMF-based approach, which offers a joint, parts-based representation of multiple modalities. Our method builds on the efficient unimodal NMF scheme proposed in (16) which implements the accelerated hierarchical alternating least squares (A-HALS) algorithm. This method has been recently developed for the joint factorisation of multi-omics data in (intNMF (17)). Here, we adapt the method to jointly factorise spatial gene expression and morphological image features derived from a matched H&E image.

The joint factorisation problem for spot-wise spatial gene expression and paired spot-wise morphological image features data is formulated (as in intNMF) as follows:

Given two non-negative input matrices *Y*_rna_ ∈ ℝ_+_^*m×n*^ and *Y*_img_ ∈ ℝ_+_^*m×f*^ where *m* is the number of spatial locations (spots), *n* is the number of genes and *f* the number of image features, building on the intNMF formulation, *CellPie* factorises the high-dimensional input matrices into a shared non-negative matrix *W* ∈ ℝ_+_^*m×k*^ and two modality specific non-negative matrices *H*_rna_ ∈ ℝ_+_^*k×n*^, where *k* ≪ *n, f* is the number of factors (rank), and *H*_img_ ∈ ℝ_+_^*k×f*^ matrices, such that:

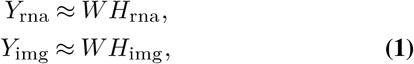

for gene expression counts and image feature data respectively.

The factors *W*, *H*_rna_ and *H*_img_ are calculated by solving the following joint optimisation problem:

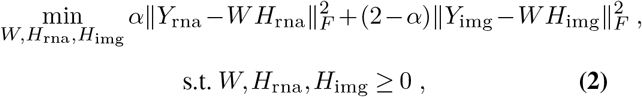

where the parameter *α* is a modality specific weight (default is set to 1.0 so that each modality is equally weighted) and ∥·∥_*F*_ is the Frobenius norm. The above non-convex optimisation problem is reduced to an alternating pair of convex optimisations through iterative updates. The accelerated hierarchical least squares (A-HALS) is adapted to jointly factorise two matrices, see (17) for details.

The image-based morphological features (*Y*_img_) are extracted from square patches centered at the spot locations of the corresponding H&E image. Features are extracted from a range of different patch sizes in order to include features at multiple scales. For example, for Visium 10x datasets, where there is a gap of 45*um* between two neighbouring spots where spatial gene expression is not measured, the default patch size is set to a range between (0.1 − 2.0) of the spot diameter in order to include morphological information that compensates for the discontinuity of the gene expression between the spots. For coarser resolution, such as the Spatial Transcriptomics where the gap between the two spots is even larger, a larger range can be used. By default *CellPie* uses spot-level pixel intensity features (bin-counts) of each color channel in the H&E image, using the histogram function of *Squidpy* (18). However, the user can import any relevant non-negative spot-wise image features, such as features extracted using deep learning architectures. An overview of *CellPie* approach is shown in Fig. 1.

**Fig. 1.**
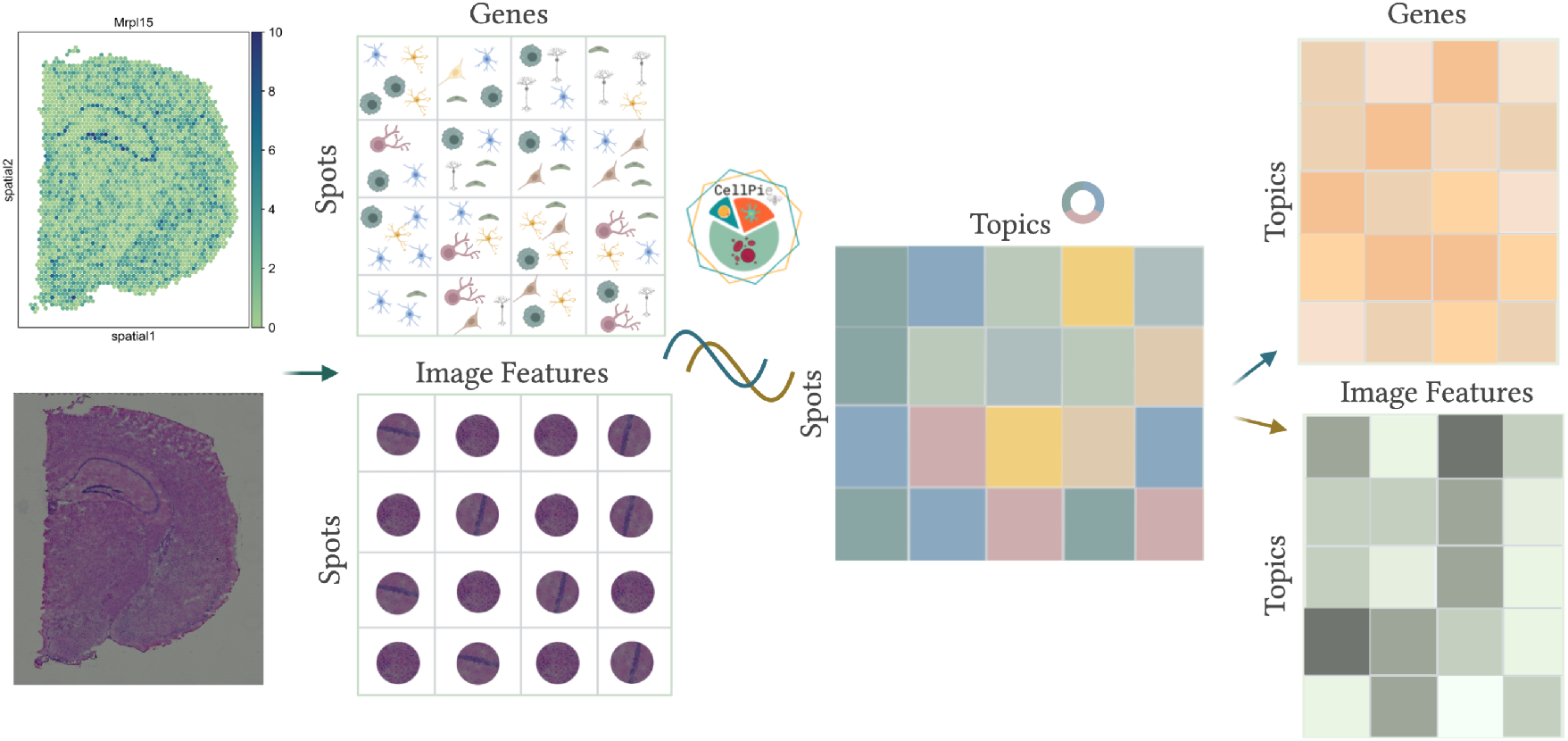
Graphical overview of *CellPie* method. *CellPie* takes as input spatial gene expression counts (spots by genes) and paired morphological image features extracted from Haematoxylin and Eosin images (spots by features). These two modalities are jointly factorised using joint non-negative matrix factorisation, resulting to three matrices: a shared spots by factors matrix, containing the reduced parts-based representation (factors) and two individual matrices containing the weights of each of the features, a gene loading and an image loading matrix.

### Model Selection and Initialisation

To help the user selecting the optimal number of factors, *k, CellPie* implements model selection through the Bi-cross validation method proposed in (19). In this method, the gene expression matrix (*Y*_rna_) are partitioned into four random blocks, as follows:

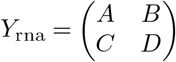

where *A* ∈ ℝ_+_^*r×s*^, *B* ∈ ℝ_+_^*r×*(*n*−*s*)^, *C* ∈ ℝ_+_^(*m*−*r*)*×s*^ and *D* ∈ ℝ_+_^(*m*−*r*)*×*(*n*−*s*)^. Blocks *B, C* and *D* are then used for training while the upper-left block *A* is predicted from *D* using the off-diagonal *B* and *C* blocks. We run *CellPie* for a range of *k* values and the optimal model is considered the one with the lowest reconstruction error. However, it is worth pointing out that prior biological knowledge of the tissue is always advantageous and *k* can also be selected in conjunction with this prior knowledge where available.

The intNMF algorithm is iterative and can converge to different locally optimal solutions, so that a good initialisation of the *W*, *H*_rna_ and *H*_img_ factors is important. *CellPie* by default uses the Non-negative Double Singular Value Decomposition (NNDSVD), an effective, non-random initialisation strategy, proposed in (20). This algorithm uses two Singular Value Decomposition (SVD) processes (we used an implementation for sparse matrices) and has been demonstrated to rapidly reduce the approximation error. Random initialisation methods are also implemented in *CellPie* however, these methods usually require multiple restarts to ensure that the algorithm converges to a reasonable solution.

### Benchmark and application to spatial transcriptomics datasets

We benchmark *CellPie* in two different human cancer types of different spatial resolution. In the first instance, we use a publicly available Visium human prostate adenocarcinoma with invasive carcinoma (21) in which pathologists annotations are provided. The second validation considers a published Spatial Transcriptomics dataset of a HER2-positive human breast cancer (22), where pathologists annotations of the tissue are also available. We evaluate the clustering performance of *CellPie* and other published dimensionality-reduction methods (*NSF, HNSF, sklearn-NMF, MEFISTO, PNMF, FA*) against the pathologist’s ground truth labels. In addition, we apply *CellPie* on a recent highly resolved spatial transcriptomics Visium HD human colorectal cancer dataset (8) to demonstrate the scalability of the algorithm for very high dimensional datasets. More information on parameter settings for competing methods is available on the supplementary material.

## Results

### Visium human prostate invasive carcinoma

Initially, we applied and benchmarked *CellPie* on a 10x Genomics Visium human prostate adenocarcinoma with invasive carcinoma dataset (Fig. 2A). 10x Genomics supplies a paired H&E image that has been annotated by a pathologist. To achieve finer granularity of the invasive carcinoma region, we re-annotated the image with more fine grained pathologist annotations of the different tissue regions (Connective tissue, Gleason 3, Gleason 4, Immune cells, Neural, Normal glands, PIN and Vascular) as shown in Fig. 2B. Consequently, the tumour region was annotated using the Gleason scoring system, labelled as Gleason 3 and Gleason 4. This enables us to evaluate if *CellPie* can identify factors that correspond to these specific Gleason areas. In prostate, two main lineages that are biologically and architecturally different can be seen: the stromal compartment and the epithelial compartment. The stroma is composed of fibroblasts, smooth muscle, nerves, and blood vessels in diverse proportions (23, 24). Prostate carcinoma arises from the glandular epithelial compartment and, unlike other organs, is not graded by individual cells differentiation but mainly by its architectural features using the Gleason grading, from 1 to 5. Due to poor reproducibility and lack of biological support, Gleason 1 and 2 are no longer reported. Gleason 3 cancers comprise the most differentiated adenocarcinoma, consisting of discrete glandular units with varying sizes and shapes (25). Individual tumor acini have smooth, typically circular edges and intact basement membranes. In contrast, Gleason 4 cancers are composed of poorly formed glandular units with indistinct borders, fused glands, and irregularly infiltrating stroma (25, 26). Intuitively, Gleason 4 tumors exhibit higher degrees of differentiation, increased cancer progression, and therefore stronger prognostic correlations.

**Fig. 2.**
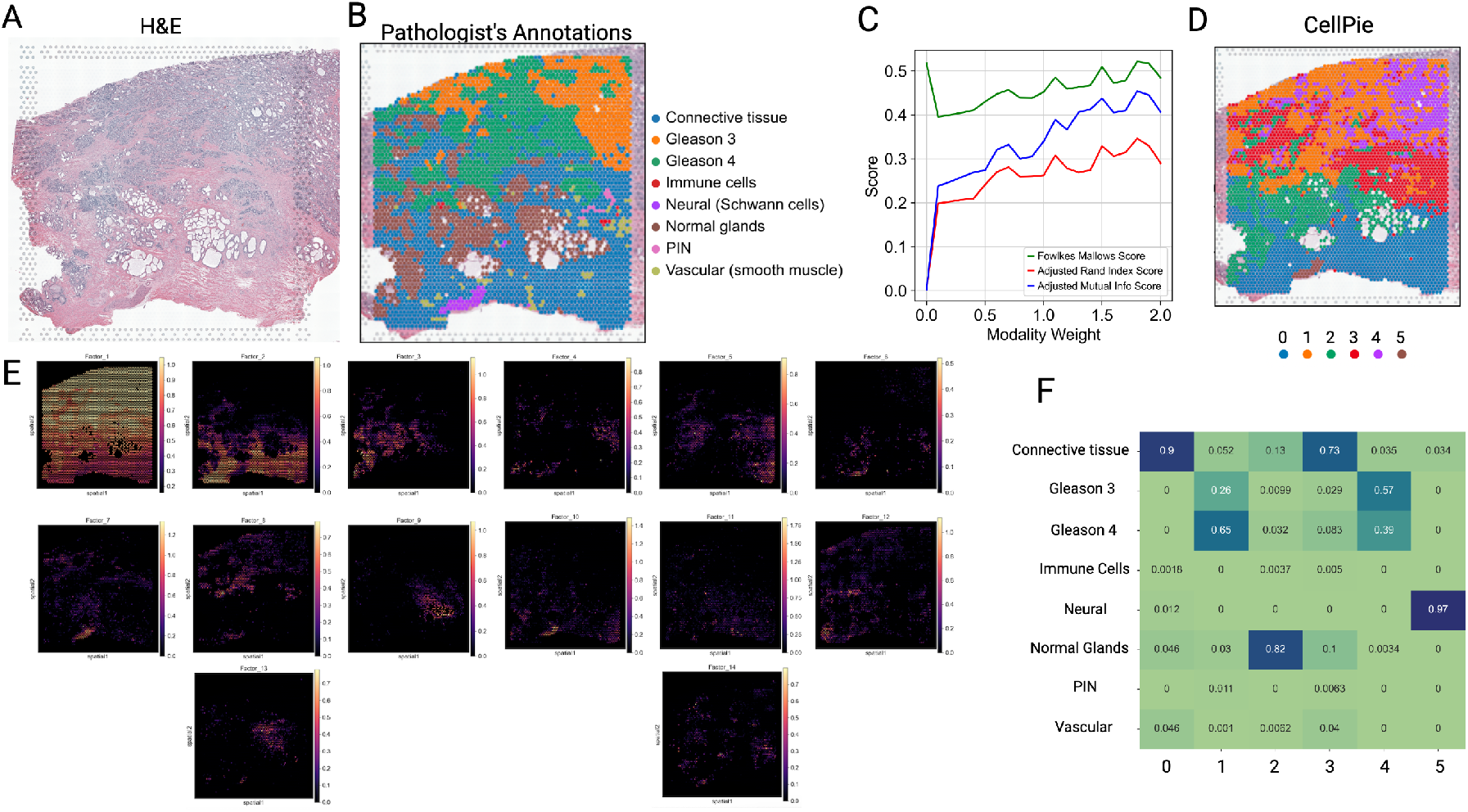
Validation of *CellPie* on human prostate cancer data, (A) Haematoxylin and Eosin image of the invasive prostate carcinoma tissue. (B) Pathologist’s annotations of the tissue. (C) Clustering performance of *CellPie* for a range of modality weights. The plot shows three measures (Fowlkes Mallows, Adjusted Rand Index and Adjusted Mutual Info) of clustering similarity between *CellPie* and the pathologist’s annotations. (D)*CellPie* clusters computed using the *leiden* algorithm on *CellPie*’s output. (E) *CellPie*’s factors. F Contingency table between *CellPie*’s clusters and pathologist’s annotations.

Model selection showed that the optimal number of factors is 14 (Supplementary Fig. 1A) and therefore we used this number for *CellPie* and all the other methods. To create labels, we clustered *CellPie*’s factors using the *leiden* clustering algorithm, where we adjusted the number of neighbours and the resolution parameters so that we ended up we 6 clusters (corresponding to 6 major pathologist’s annotated areas, since PIN and Vascular are only found in a few spots). First, we explored how the modality weight affects the *CellPie*’s clustering performance by running *CellPie* across a range of modality weights spanning from 0.0 (image only) to 2.0 (RNA expression only). We then computed the Fowlkes Mallow, Adjusted Rand Index and Adjusted Mutual Info scores between the *CellPie*’s clusters and pathologist’s annotations, as shown in Fig. 2C. The optimal modality weight was found to be 1.8, indicating that the integration of the two modalities leads to a better clustering performance than only using the expression modality (weight= 2.0). *CellPie*’s factors and their clustering are shown in Fig. 2D and Fig. 2E. To label the clusters, we computed the cross tabulation between *CellPie*’s clusters and the pathologists annotations, where clusters 0 and 3 correspond mainly to the connective tissue region, cluster 1 to Gleason 4 and parts of Gleason 3, while cluster 2 aligns with the normal glands. Cluster 4 is a mix of Gleason 3 and Gleason 4 and cluster 5 corresponds to the neural region (Fig. 2F). Overall *CellPie* can identify the 6 major tissue areas with a corresponding ARI of 0.41, showing good agreement with the ground-truth (the fact that the connective tissue is split into two clusters results to a lower ARI ((Fig. 2F))).

We then benchmarked *CellPie, NSF, NSFH, MEFISTO, FA, PNMF* and *CellPie* with only gene expression (this corresponds to standard single NMF) against the pathologist’s ground-truth. Similarly to *CellPie*, we ran each method according to author’s default settings with 14 factors and then we clustered the factors using *leiden* clustering, while tuning the number of neighbours and resolution parameters to result into 6 clusters (Fig. 3A). The ARI between the ground-truth and each method is shown in fig. 3B, where *Cellpie* shows the highest performance. In addition, *CellPie* is the only method that can identify the two separate tumour region areas, Gleason 3 and Gleason 4.

**Fig. 3.**
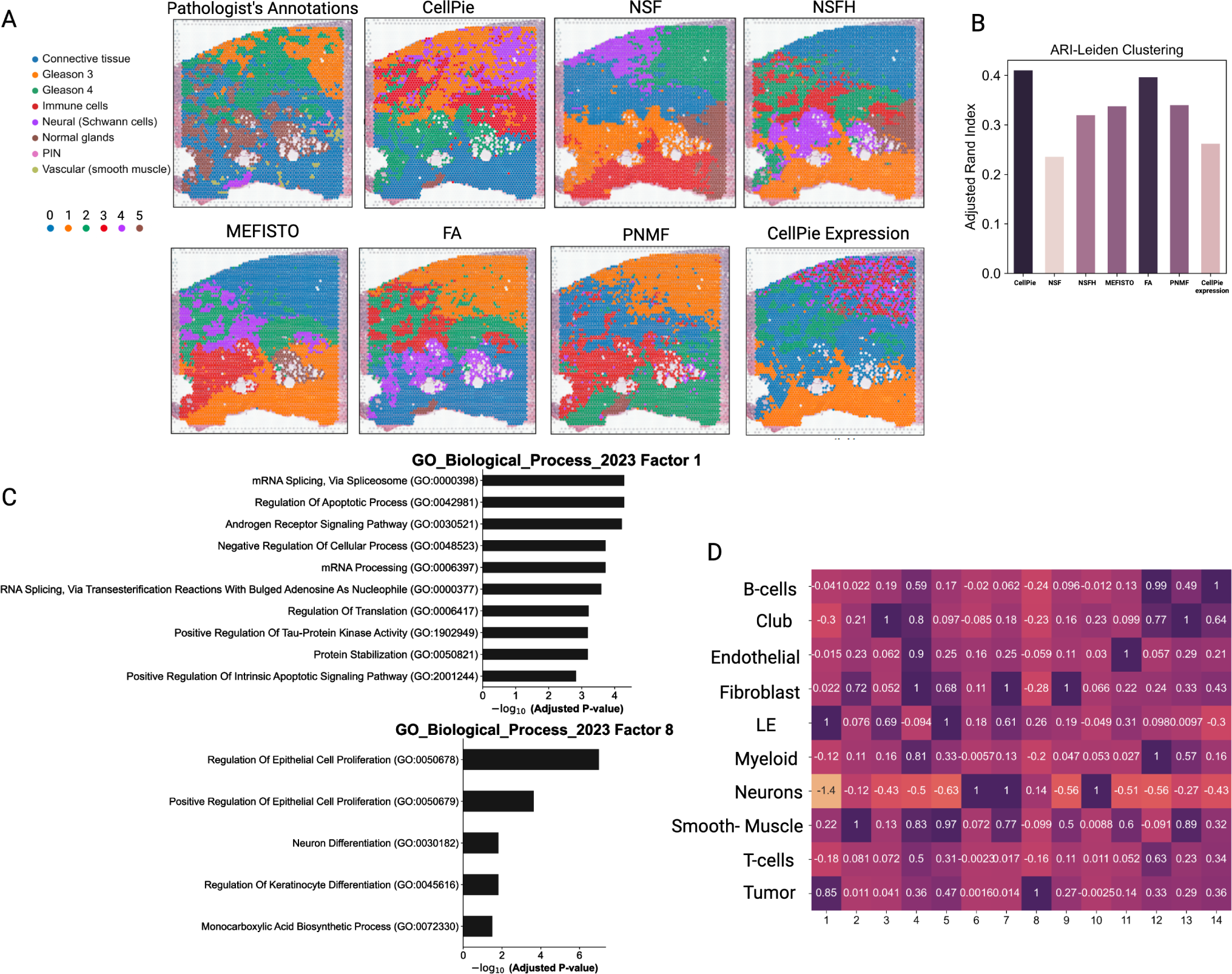
Comparison of *CellPie* against other published dimensionality reduction methods: (A) Pathologist’s annotations and clustering results of *CellPie, NSF, NSFH, MEFISTO, FA, PNMF* and *CellPie* with only gene expression. To cluster the factors of each method, the *leiden* clustering algorithm with 6 clusters was used. (B) Adjusted Rand Index between the pathologist’s annotations and the clusters of the methods discussed in (A). (C) GO ontology of Factor 1, using *CellPie*’s top 150 marker genes of *CellPie*’s Factor 1. (D) Highest ranked cell-types per factor, computed using a published single-cell RNA-seq dataset and Scanpy’s score_genes function.

To further annotate the factors with cell-types, we interrogate *CellPie*’s gene loading matrix in conjunction with a modified cell signature obtained from (27), with the addition of markers for B-cells (28) and neurons (using scanpy.tl.score_genes function). In Fig. 3D, the factor interpretation of the invasive prostate carcinoma is shown. Factors 1, 3 and 5 were in keeping with the epithelia compartment (luminal epithelial - LE), based on their score. Factor 1 represents the glandular area of the prostate covering both benign and malignant (tumour) area, with high expression of *KLK2, KLK3* and *ACPP* (Supplementary Fig. 2), genes associated with prostate glandular epithelium marker. Amongst top genes in Factor 3, *CellPie* identified *MSMB* and *AZGP1* which are usually expressed in benign glandular tissue. Factors 2, 4, 5, 7 and 13 correspond to the stromal compartment (smooth-muscle) where there is high expression of *MYL9, ACTG2, ACTA2* and *TAGLN*, all involved in muscle contraction. Tumour cells are mainly found in Factors 1, 8 and in some parts of Factors 4 and 5 (Fig. 3D). To further validate the biological significance of the regions identified by our method at the molecular level, we applied Gene Ontology (GO) analysis (Fig. 3C). The GO terms most enriched in Factor 1, which shows the tumor-specific expression pattern, include Androgen Receptor signaling pathway and mRNA alternative splicing. The former is the main driver of the progression of prostate cancer, while the latter has been reported in recent years to play a significant role in prostate cancer (29). The enrichment of apoptosis-related entries may reflect the inhibition of apoptotic signaling in prostate cancer (30).

*CellPie* has demonstrated a particular advantage in prostate cancer samples by identifying a Gleason 4 specific factor, Factor 8. The two main GO terms enriched by this factor are “Regulation of Epithelial Cell Proliferation” and “Positive Regulation of Epithelial Cell”, indicating that the cancer cells at this stage have entered a rapid proliferation phase. Additionally, “Proliferation Regulation Of Keratinocyte Differentiation” may reflect changes in integrin and further alterations in the microenvironment, suggesting enhanced invasive capacity (31). As prostate cancer progresses, significant metabolic pathway changes occur, including carbohydrate metabolism, lipid metabolism, and alterations in 1C metabolic homeostasis, processes highly related to the “Monocarboxylic Acid Biosynthetic Process” (32). Interestingly, Factor 8 also includes the term “Neuron Differentiation”, and a moderate signal can be observed in Schwann cells in the image, possibly indicating a functional relationship between Gleason 4 and Schwann cells. This finding aligns with previous reports that Schwann cells in prostate cancer increase integrin-dependent tumor invasion on laminin (33).

Analysis of the top genes within Factor 8 reveals that most of these genes have already been reported to be involved in the development of prostate cancer. For example, the genes *CRISP3, HPN*, and *AMACR* are significantly upregulated in prostate cancer and can serve as biomarkers (34–36). More importantly, some top genes such as *GDF15* are involved in the TGF-beta signaling pathway, which has been reported to be involved in prostate cancer metastasis (37, 38), suggesting an increased invasive and metastatic capability of tumor cells at this stage. The *ERG* gene, which participates in the Wnt/*β*-catenin signaling pathway, is highly expressed in malignant prostate cancer and promotes tumour progression (39). High *ERG* expression is also associated with a high Gleason score (39).

Factor 10 may be the Schwann cell-specific factor. Based on the images, Factor 10 shows the best consistency with the regions corresponding to Schwann cells. This is consistent with the enrichment analysis results. The GO terms enriched in Factor 10 are primarily related to nervous system development and myelination. Analysis of the top genes in factor 10 reveals some genes specifically expressed in the myelin sheath. For example, the calcium-binding protein *S100B* is a common marker for Schwann cells, and the *MPZ* gene is specifically expressed in Schwann cells, being a major structural protein in the myelin sheath (40).

Factor 2 exhibits high expression in normal connective tissue and has enriched GO terms that include “Homotypic cell-cell adhesion”, “Supramolecular Fiber Organization” and “Integrin-Mediated Signaling Pathway”. These terms highlight smooth muscle and collagen fibre intra and extracellular organization, which are key components of the connective tissue found in the prostate.

### HER2-Positive Breast Cancer

For the second validation, we used a published HER2 (human epidermal growth factor receptor 2)-positive breast cancer Spatial Transcriptomics (ST ≈ 0 − 200 cells/spot) dataset (patient H1) (22), where pathologist annotations are available and were used as ground-truth (Fig. 4A and 4B). The regions identified by the pathologists were labeled as: adipose tissue, breast glands, cancer in situ, connective tissue, immune infiltrate and invasive cancer. Model selection was performed for 3 repetitions and the two resulted in 4 optimal factors while the third in 5 factors (Supplementary Fig. 1B). We used 5 because this number is closer to the number of ground-truth labels. We used this number for *CellPie* and all the other methods. To select the optimal modality weight, we fit *CellPie* for a range of weights between 0.0 and 2.0 and we selected the one that corresponds to the best ARI between the pathologist’s annotations and *CellPie*’s clustering (Supplementary Fig. 1C). The resulting factors are shown in Fig. 4B.

**Fig. 4.**
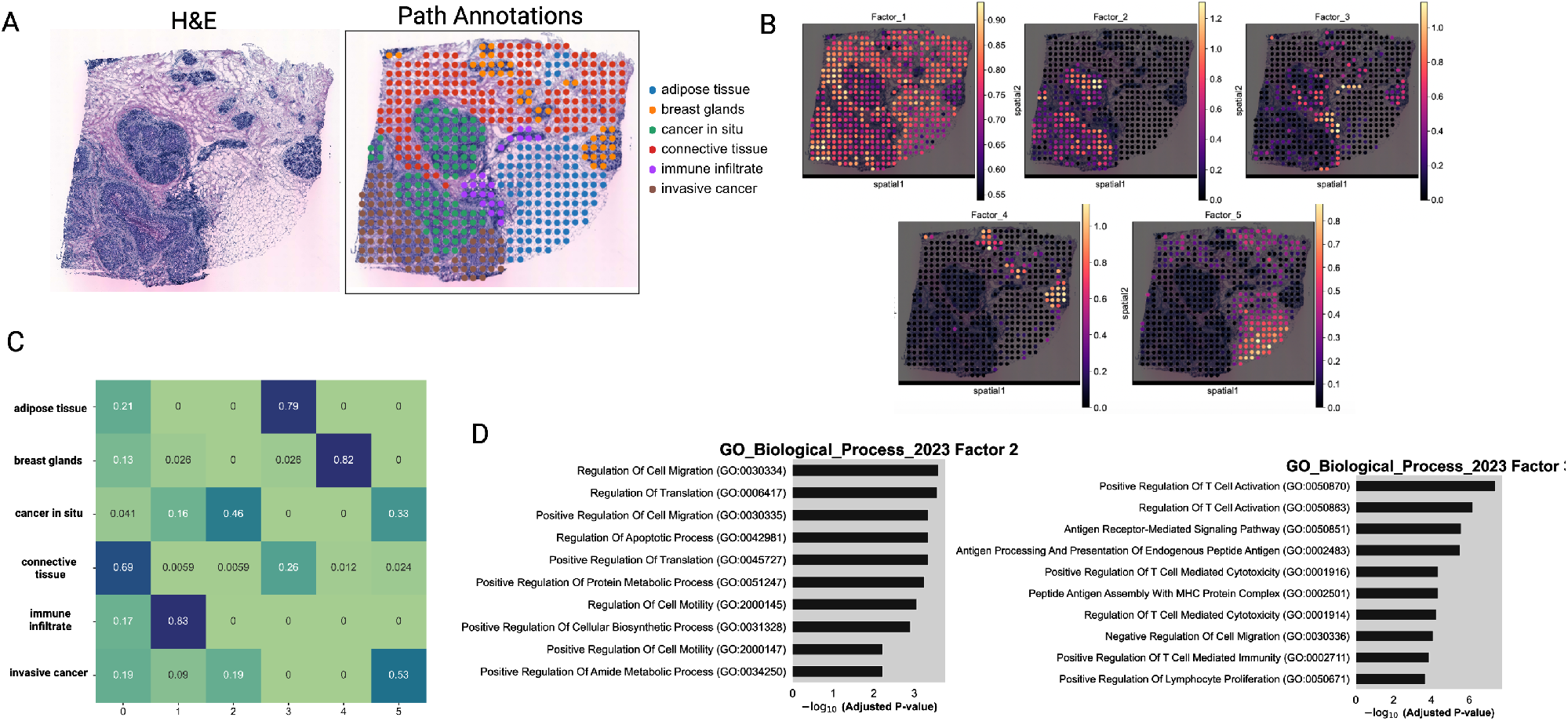
(A) H&E image of the HER2-positive Breast Cancer sample (Patient H1) and pathologist’s annotations. (B)*CellPie*’s factors. (C) Contingency matrix between *CellPie*’s factors and pathologist’s annotations. (D) GO ontology on Factors 2 and 3.

To compare the resulting factors of each method with the ground-truth pathologist’s annotations we clustered the factors using *k-means* algorithm with *k* = 6 clusters, so that it reflects the number of the ground-truth labels (Fig. 5A). *CellPie* and *NSF* show the best concordance with the pathologist’s annotations (Fig. 5B), distinguishing breast glands, immune infiltrates, adipose tissue with good accuracy (Fig. 4C) while connective tissue, cancer in situ and invasive cancer regions are a mix of 2 clusters.

**Fig. 5.**
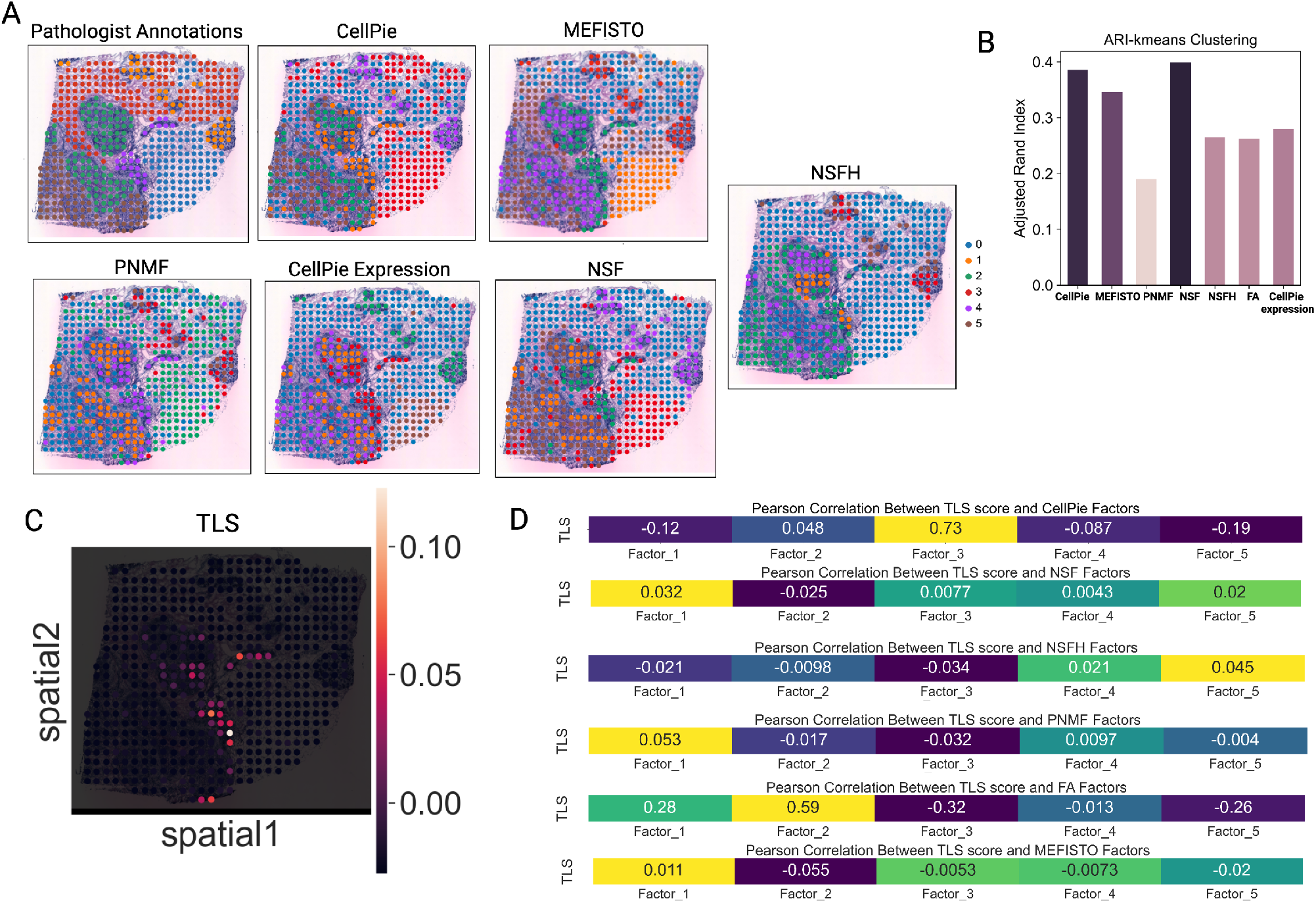
(A)Pathologist’s annotations and clustering results of *CellPie, MEFISTO, PNMF, CellPie* with only gene expression, *NSF* and *NSFH* factors. To cluster the factors of each method, the *k-means* clustering algorithm with 6 clusters was used. (B) ARI between the resulting clusters and the ground-truth. (C) Spatial distribution of the tertiary lympoid structures (tls) score, computed using the results in (22). (D) Pearson correlation between *CellPie*’s, *NF* ‘s, *NSFH*’s, *PNMF* ‘s and *FA*’s factors and the tls scores.

It has been shown that in HER2-positive tumours, tertiary lymphoid structures (TLSs) have predicted values for clinical outcomes and treatment responses. These structures, which resemble lymph nodes, can form ectopically in tissues such as tumours, providing anti-tumour immune responses (22, 41). As shown in (22) TLS sites are enriched in B and T-cells. We used the TLS score results provided in (22) (Fig. 5C) to investigate if any of the resulting factors aligns with the TLS enriched areas. For each method, we computed the Pearson correlation between the TLS score and each of the factors (Fig. 5D) and we found that the largest correlation (0.73) was in *CellPie*’s Factor 3 (corresponds to immune infiltrate region).

The enrichment results of Factor 3 mainly focus on items related to T-cell-mediated immune responses. From the images, the areas corresponding to Factor 3 include immune infiltrates, cancer, and breast glands, with the strongest signal observed in the immune infiltrates. This is consistent with previous reports that breast cancer tissues contain infiltrates dominated by activated T lymphocytes (42).

Factor 2 overlaps highly with two cancer areas, cancer in situ and invasive cancer. The enriched GO terms include cell migration, apoptosis, and protein metabolic processes, which are consistent with the increased infiltration capability of cancer cells at this stage.

### Visium HD Human Colorectal Cancer

Visium HD provides single-cell or near single-cell spatial resolution, depending on the bin size, offering a comprehensive view of spatial gene expression patterns. However, due to the high dimensionality of the data (resolution at 2*um*, 8*um* and 16*um*), current factor analysis methods either require a very long time to run on a standard computer or become impractical.

We applied *CellPie* to a Visium HD human colorectal cancer (CRC) dataset from patient P1 (8). Given the key role of the immune cellular landscape in the CRC progression, in their paper, the authors studied the immune cellular composition of the tumour microenvironment of the CRC. Amongst other immune cellular populations, macrophages were found to be the most abundant in the tumour periphery. In that region, the authors identified two pro-tumour macrophage subpopulations forming distinct spatial niches. These populations are mainly defined by the expression of *SELENOP* and *SPP1* genes so the authors defined the two distict macrophage subpopulations as SELENOP+ and SPP1+. Differential gene expression showed enrichment of the *REG1A* and *TGFBI* genes for the SELENOP+ and SPP1+, respectively. Therefore, we wanted to test whether *CellPie* can identify factors that are related to these two macrophage types. We ran *CellPie* on the 16*um* bin resolution for 80 factors (Supplementary Fig. 1D, Supplementary Fig. 3) across a range of different modality weights, and computed the Pearson correlation between the resulting factors and the expression of *REG1A* and *TGFBI* genes. For all the tested weights, there is a good correlation between Factor 18 and REG1A expression, while the correlation between SPP1+ associated factor (Factor 9) and *TGFBI* was lower, although still acceptable (Fig. 6A and Fig. 6B). The best correlation found was for modality weight 1.5, highlighting the importance of integrating image and expression features. Gene ontology, using the MSigDB Hallmark gene sets, on Factor 9 (SPP1+), showed enrichment in pathways such as cholesterol homeostasis. Factor 18 (SELENOP+) showed enriched pathways such as TNF-alpha signaling via NF-kB and UV response that are similar to those found in (8) (Fig. 6C).

**Fig. 6.**
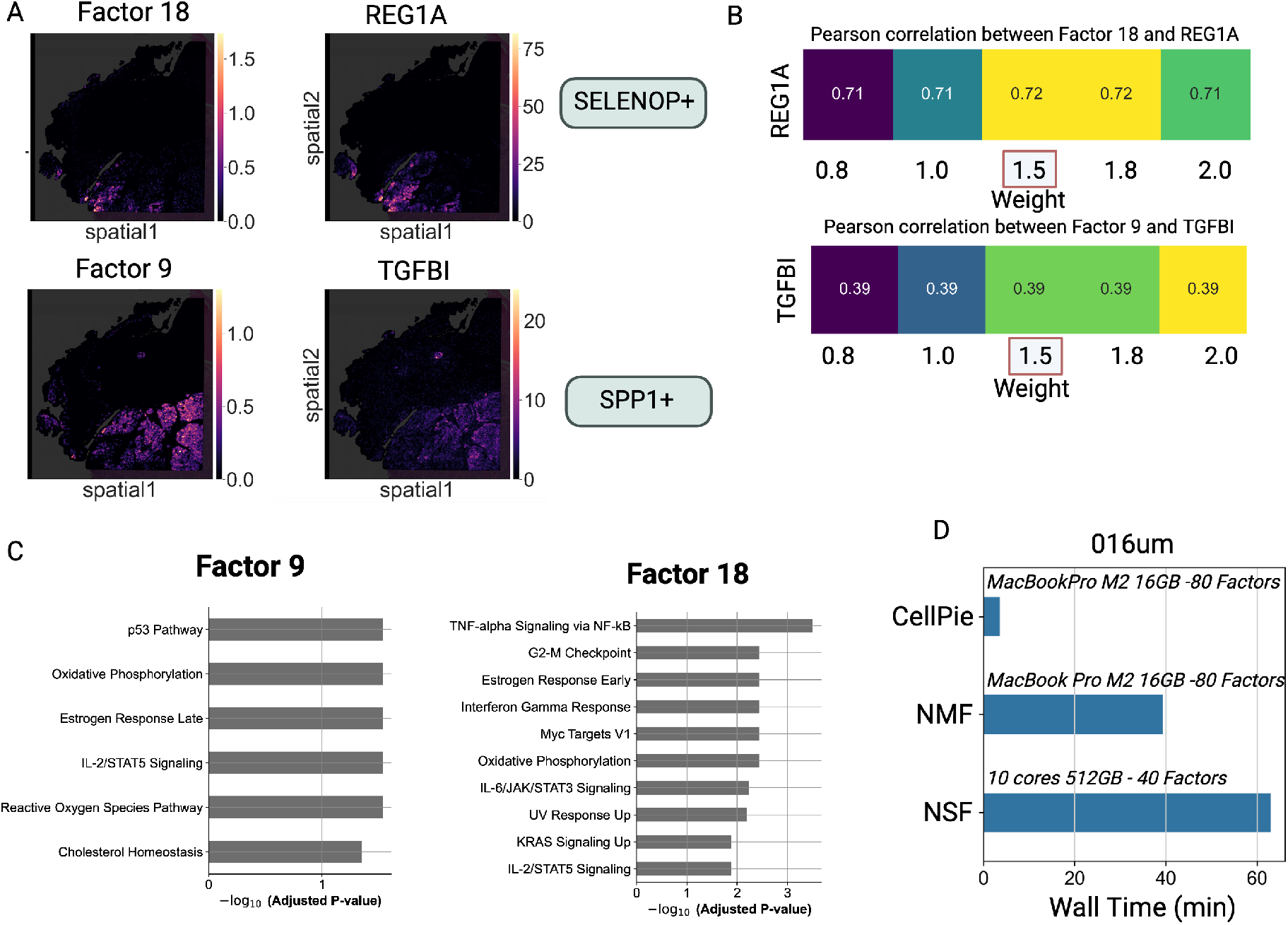
(A)Top: *CellPie*’s Factor 18 and spatial gene expression of *REG1A* gene, marker of SELENOP+ macrophages. Bottom: *CellPie*’s Factor 9 and spatial gene expression of *TGFBI* gene, marker of SPP1+ macrophages. (B)Top: Pearson correlation between *CellPie*’s Factor 18 and the *REG1A* gene for a range of weights. Bottom: The same but between Factor 9 and the *TGFBI*. (C) Pathways enriched in Factors 9 and 18, respectively, where the MSigDB Hallmark 2020 database was used. (D) Convergence time of *CellPie*, sklearn *NMF* and *NSF*.

We also investigate the convergence time of *CellPie*, sklearn’s *NMF* and *NSF*, where *CellPie* and sklearn’s *NMF* ran on a standard MacBook Pro M2 with 16*GB* memory while *NSF* failed to ran on those settings due to memory issues. *NSF* with 80 factors also failed to run on a high performance computing cluster with 10 cores and 512*GB* memory. To demonstrate the computational performance we ran *NSF* with 40 factors (Fig. 6D). For the 16*um* bin resolution, *CellPie* shows a significantly faster computational performance of about 4*min*, while *NMF* and *NSF* (we used 3, 000 inducing points for *NSF*, its complexity scales cubically with the number of inducing points) converged in about 39*min* and 63*min* respectively (Fig. 6D). For the higher resolution of 8*um* (507, 684 spots), *CellPie* converged in 6*min*43*sec*, while we were not able to ran *NSF* and *NMF* due to lack of memory.

## Conclusion

In this paper, we presented *CellPie*, a scalable and efficient factor discovery method for spatial transcriptomics that leverages a fast implementation of non-negative matrix factorisation to integrate spatial transcriptomics counts data with histopathological image features. *CellPie* significantly reduces the computation time and memory requirements in comparison to other methods, making it applicable to very high-dimensional spatial transcriptomics datasets such as those from the latest Visium HD technology.

Our evaluation on two different human cancer types and spatial resolutions demonstrated that *CellPie* is among the top-performing methods, achieving performance very close to the best method in terms of the overall clustering accuracy. Furthermore, *CellPie* outperforms existing methods in the identification of Gleason 3 and Gleason 4 areas in the invasive prostate carcinoma, where *CellPie* was the only method able to distinguish between those two types. In the HER2-positive breast cancer dataset, *CellPie* effectively identified key regions corresponding to different tissue types and showed the highest factor correlation to the tertiary lymphoid structures. Furthermore, in the Visium HD colorec-tal cancer dataset, *CellPie* identified factors associated with distinct macrophage subpopulations with specific gene expression patterns, while maintaining high computational efficiency.

The integration of histopathological features with gene expression has shown to improve the accuracy of spatial transcriptomics factor analysis. By adjusting the relative weight of the two modalities, *CellPie* provides a flexible approach to factor discovery, enabling the extraction of meaningful spatial patterns from complex tissue samples.

One limitation is that *CellPie* does not explicitly model the spatial dimension of the data, e.g. the neighbourhood relationships between spots. Hence, a spatially aware version of the joint NMF algorithm would be an interesting future direction. Furthermore, *CellPie* and the intNMF algorithm (17), which *CellPie* is based on, could be extended to jointly factorise more than two matrices simultaneously. This enhancement would allow for integration of several omics datasets possessing a shared dimension (e.g. including spatial protein markers).

## Acknowledgments

We thank Alkmini Damkou, a PhD student in Simons lab (TUM-NCB) for our fruitful discussions.

## Data availability

The *CellPie* code and the notebooks to reproduce the analysis are available as an open-source python package in https://github.com/ManchesterBioinference/CellPie. The prostate invasive carcinoma pathologist’s annotations are also available on the github repository. The Human Prostate Cancer can be downloaded from https://www.10xgenomics.com/datasets/human-prostate-cancer-adenocarcinoma-with-invasive-carcinoma-ffpe-1-standard-1-3-0, the HER2-positive can be found in https://github.com/almaan/her2st while the Visium HD colorectal cancer can be downloaded from https://www.10xgenomics.com/products/visium-hd-spatial-gene-expression/dataset-human-crc.

