## Supplementary material for "*CellPie*: a scalable spatial transcriptomics factor discovery method via joint non-negative matrix factorization"

### 1 Parameter settings

For the spatial transcriptomics data, we used normalised and logarithmised counts for *CellPie*, using Scanpy's normalize_total and log1p functions. Same normalisation was used for *MEFISTO*, *FA* and *NMF*. Image features where logarithmised in the same way as the gene expression data. We note that for the Visium HD datasets we used raw gene expression counts. We used raw counts for *NSF*,*NSFH* and *PNMF*. As in *NSF* paper, we used the top $2,000$ informative genes using Poisson deviance for *NSF*,*NSFH*,*PNMF* and *MEFISTO*. We ran *NSF* and *NSFH* using $3,000$ inducing points with Poisson likelihood for prostate cancer and crc datasets and 500 inducing points for the HER2-positive breast cancer dataset. For *MEFISTO* we used Gaussian likelihood with $1,000$ inducing points for the Visium and Visium HD datasets and 500 for the ST dataset. Scripts to reproduce the results are available in `https://github.com/ManchesterBioinference/CellPie`. For computational time comparison purposes only, we ran *NSF* on the Visium HD CRC dataset with 40 factors, using raw counts. As the 'preprocess.deviancePoisson' failed for that dataset, we used all the genes in the dataset and we filter only those which are expressed in at least one spot and with at least 100 counts across all the spots.

### 2 Supplementary Figures

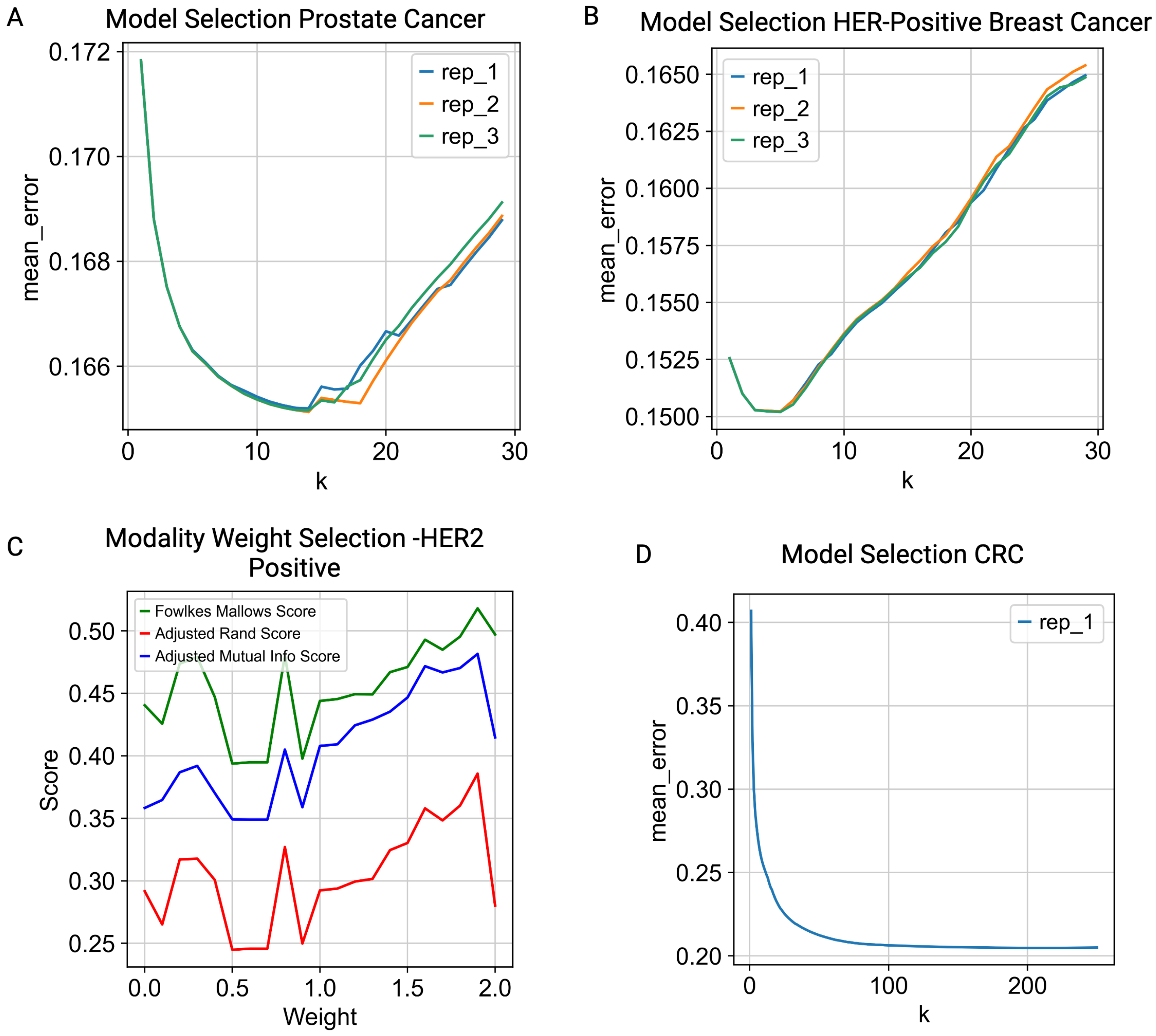


Figure 1: (A) Model selection for the human prostate adenocarcinoma with invasive carcinoma dataset. (B) Model selection for the HER2-positive breast cancer dataset. (C) Modality weight selection for human HER2-positive breast cancer.(D) Model selection for the human colorectal cancer dataset.

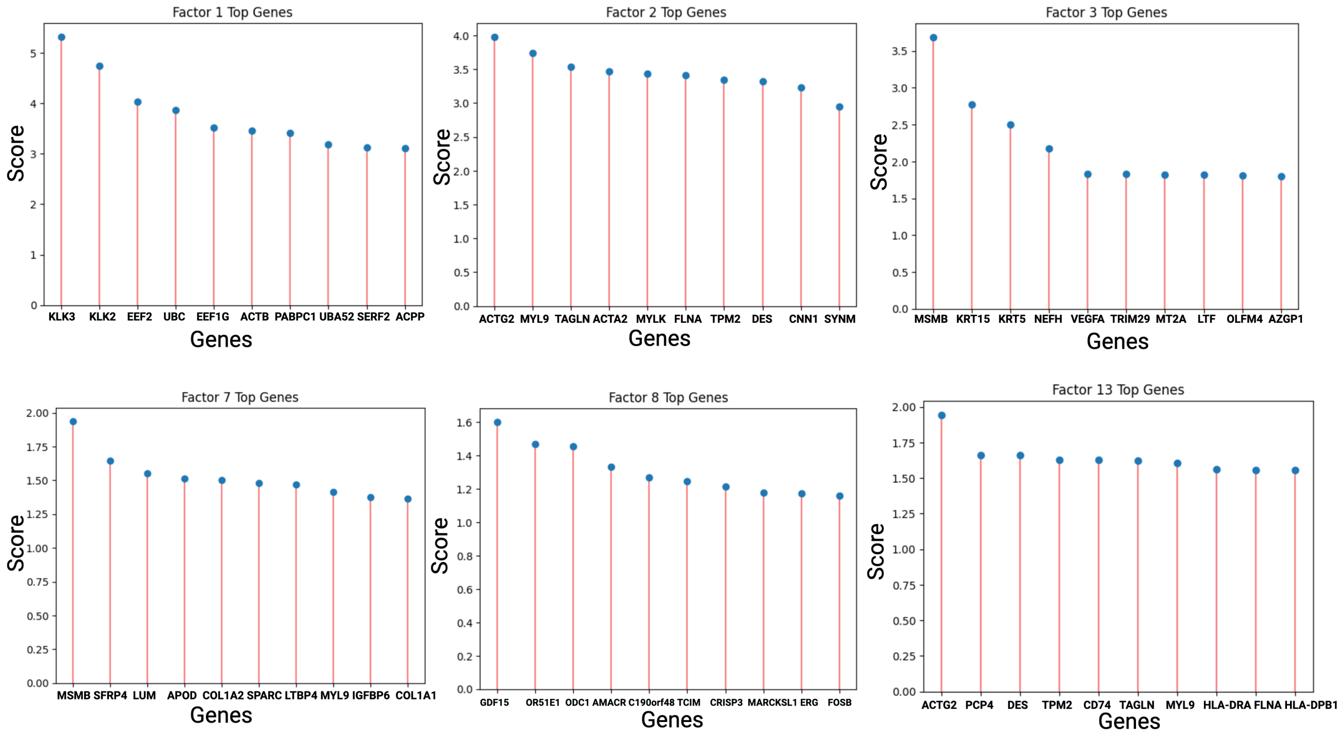


Figure 2: Plots showing the score of the top 10 marker genes of Factors 1,2,3,7,8 and 13.
